# *Kcnn3* as a target for treating aberrant behaviors in stressed, ethanol-dependent mice

**DOI:** 10.1101/734970

**Authors:** Audrey E. Padula, Jennifer A. Rinker, Fauzan Khan, Marcelo F. Lopez, Megan K. Mulligan, Robert W. Williams, Howard C. Becker, Patrick J. Mulholland

**Author notes:** Corresponding Author: Patrick J. Mulholland, PhD.

## Abstract

Anxiety and mood disorders are often comorbid with alcohol use disorder (AUD) and are considered critical in the development, maintenance, and reinstatement of alcohol dependence and harmful alcohol-seeking behaviors. Because of this high comorbidity, it is necessary to determine shared and unique genetic factors driving heavy ethanol drinking and anxiety-related behaviors. We used a model of stress-induced escalation of drinking in ethanol dependent C57BL/6J mice to measure anxiety-like behaviors on the marble burying and novelty-suppressed feeding task (NSFT) during abstinence. In order to identify novel pharmacogenetic targets that may lead to more effective treatment, a targeted bioinformatics analysis was used to quantify the expression of K^+^ channel genes in the amygdala that covary with anxiety-related phenotypes in the well phenotyped and fully sequenced family of BXD strains. A pharmacological approach was used to validate the key bioinformatics finding in ethanol-dependent, stressed C57BL/6J mice during the NSFT. Amygdalar expression of *Kcnn3* correlated significantly with just over 40 anxiety-associated phenotypes. Further examination of *Kcnn3* expression revealed a strong eigentrait for anxiety-like behaviors in this family. *Kcnn3* expression in the amygdala correlated negatively with binge-like and voluntary ethanol drinking. C57BL/6J mice treated with chronic intermittent ethanol exposure and repeated swim stress consumed more ethanol in their home cages and showed hypophagia on the NSFT during prolonged abstinence. Pharmacologically targeting *KCNN3* protein with the K_Ca_2 channel positive modulator 1-EBIO decreased ethanol drinking and reduced latency to approach food during the NSFT in ethanol-dependent, stressed mice. Collectively these validation studies provide central nervous system mechanistic links into to the covariance of stress, anxiety, and AUD in the BXD strains. Further this analytical approach is effective in defining targets for treating alcohol dependence and comorbid mood and anxiety disorders.

## Introduction

Alcohol (ethanol) use disorder (AUD) is a devastating disruption of brain function and behavior. While environmental factors and exposure are obviously critical, heritability estimates of AUD in many modern societies are high—60% or greater. Defining and understanding genetic variants among individuals in regards to AUD resistance, susceptibility, and relapse risk will provide key data to develop more personalized prevention and treatment plans (1, 2). AUD is frequently comorbid with affective mood disorders (3, 4) and this fact often limits available treatment interventions (5). The negative affective disturbances that tend to develop during abstinence from drinking can drive relapse-like behaviors in both humans and rodent models (6–12). This is further complicated by preclinical and clinical evidence that links increased alcohol drinking with the use of drugs such as SSRIs that are commonly used to treat depression and anxiety (13, 14). Moreover, alcohol consumption increases depressive symptoms and the side effects of antidepressants. Because of these limitations and the fact that current FDA-approved pharmacotherapies for AUD do not target mood disorders, it is essential to identify new targets that treat both excessive alcohol drinking and negative affective states. Despite increased efforts to identify mechanisms that drive excessive drinking and negative affective disturbances, treatment options for individuals with AUD are still limited and moving toward a personalized medicine approach.

Potassium (K^+^) channels are a large, diverse class of ion channels that modulate many aspects of the biophysical membrane properties of neurons and glia and ultimately influencing neural circuitry. Recently, a number of K^+^ channels have been implicated in neuropsychiatric conditions, including AUD and mood disorders. For example, chronic ethanol exposure produced adaptations in K^+^ channels that influenced intrinsic excitability and excessive intake (15). Pharmacological targeting of K^+^ channels reduces ethanol self-administration and intake in rodents with a heavy drinking phenotype (2, 15). Knockdown of *Kcnk9* in monoaminergic neurons (serotonergic and noradrenergic) reduces negative affective behaviors in mice (16). In a social defeat model of depression, mice that were resilient to depressive-like behaviors had higher expression of *Kcnq3* in the ventral tegmental area (VTA) and increased K^+^ channel currents in VTA dopaminergic (DA) neurons (17, 18). Friedman and colleagues (19) demonstrated that retigabine—a Kv7 channel positive modulator—restored normal firing patterns of VTA neurons and prevented the development of depression-associated behaviors. In addition, these authors reported that overexpression of *Kcnq* channels in VTA dopaminergic neurons normalized neuronal hyperactivity and depression-like behaviors in susceptible mice. In rats, Kang and colleagues confirmed that retigabine microinfusion into the lateral habenula reduced anxiety-like behaviors during alcohol withdrawal (20). Additional K^+^ channels, such as K_Ca_2, GIRK1, and Kv1.3, influence anxiety-related behaviors (21–23). These promising findings have motivated our genetic studies on the role of K^+^ channels as a factor linking negative affective behaviors and excessive drinking.

Stressful life events can precipitate the onset of major depressive disorder and are a major risk factor for developing mood disorders (24, 25), and the relationship between stress and mood disorders is influenced by gene-environment interactions (26, 27). In this study, we evaluated the extent to which chronic ethanol and stress co-exposure produces excessive drinking (28–30) and cognitive impairments (31), as well as the covariance to affective and anxiety-like behaviors. Next, we used a deep phenome and gene expression data set acquired over the past decade for many members of the BXD family of mice (32) to identify K^+^ channel genes in the basolateral amygdala (BLA) that are associated with anxiety-like behaviors. BXD RI strains are generated by intercrossing C57BL/6J and DBA/2J inbred parental strains and their isogenic progeny to generate recombinant strains that are then inbred to produce stable RI lines (33). The BXD panel has been fully genotyped and sequenced, expression profiling has been completed for numerous brain regions, and over 7,500 behavioral, morphological, and pharmacological traits have been measured across this population. Thus, BXD RI strains are well suited for systems genetics analysis of behavioral traits and are a powerful resource to study the genetic diversity of anxiety- and alcohol-related phenotypes (34–37). Once gene targets were identified, we determined if expression of top targets such as *Kcnn3* covary with voluntary ethanol drinking. Because we identified a promising candidate with correlations between anxiety-like behaviors and ethanol consumption, we used a direct pharmacological approach to validate a key role of KCNN3 protein (Kea2 channels) in jointly influencing chronic ethanol-stress, ethanol drinking, and anxiety.

## Materials and Methods

### Animals

Adult, male C57BL/6J mice (Jackson Laboratory, Bar Harbor, ME) were 8–10 weeks of age at the start of each experiment and were individually housed in a temperature- and humidity-controlled vivarium. Mice were kept on a 12 h reverse light/dark cycle with *ad libitum* access to food (Harlan Teklad Diet 2918) and water with the exception of when mice were tested on the novelty suppressed feeding test, as described below. All mice were treated in accordance with the NIH Guide for the Care and Use of Laboratory Animals (8th edition, National Research Council, 2011) under protocols approved by the Institutional Animal Care and Use Committee at the Medical University of South Carolina.

### Ethanol drinking and dependence

To establish baseline drinking, mice consumed ethanol in their home cage using a standard two-bottle choice (15% ethanol (v/v) vs water) limited-access (2 h) protocol for 4-6 weeks. Mice were pseudorandomly divided into air or ethanol vapor groups, and half of the mice then underwent two repeated weekly cycles of chronic intermittent ethanol (CIE) exposure or air (control group) in vapor inhalation chambers, alternated with two weekly home cage drinking sessions (Becker and Lopez, 2004; Padula et al, 2015; Lopez and Becker, 2005). Air or CIE exposure was delivered 16 h/day for four consecutive days followed by 72 h of abstinence prior to limited access to water or ethanol in their home cage for five test drinking days. There was 72 h between the last test drinking session and the start of vapor exposure. Body weights were recorded weekly during drinking weeks and daily during cycles of CIE exposure. The full behavioral model is presented in **Fig 1A**.

**Figure 1.**
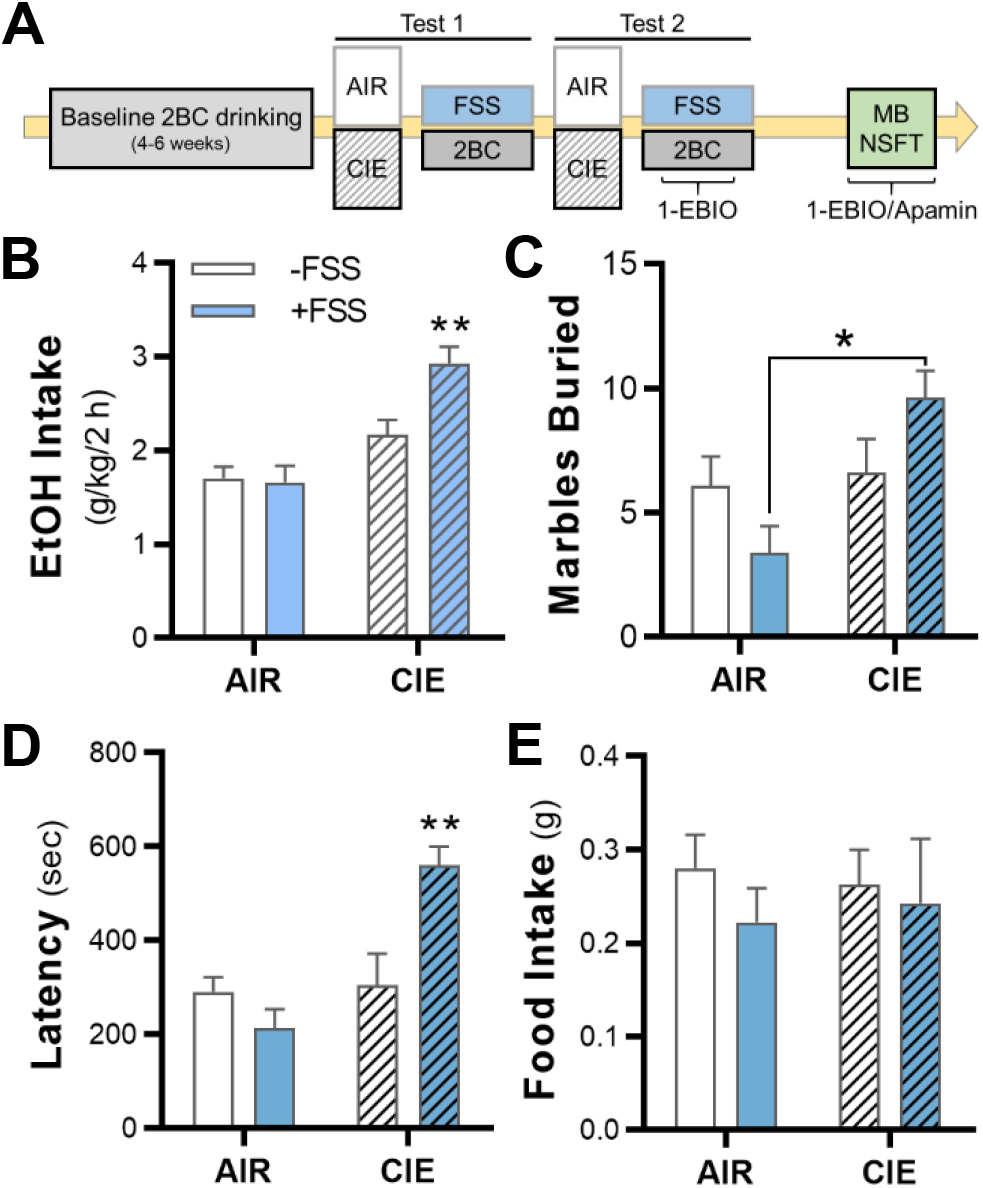
Drinking and anxiety-like behaviors in mice co-exposed to forced swim stress (FSS) and chronic intermittent ethanol (CIE). **(A)** Schematic of two-bottle choice drinking in mice exposed to CIE and FSS that were tested on the marble burying and novelty-suppressed feeding tests (NSFT). 1-EBIO or apamin were administered 30 min prior to home cage drinking or the NSFT. **(B)** Ethanol consumption in mice exposed to CIE and daily FSS 4 h prior to home cage ethanol drinking. **(C)** The number of marbles buried in a 30-min test period. **(D,E)** Latency to consume food on the NSFT and post-test food consumption in the home cage.

### Forced Swim Stress

Half of the air and CIE exposed mice were subjected to a 10-min forced swim stress (FSS) test in a glass cylinder (20 cm diameter × 40 cm high) that was half-filled with 23-25°C tap water following previously reported methods (28–30). After removal from the swim test cylinders, mice were hand-dried and placed on a heating pad for 5-10 min. For mice in the home cage drinking studies, mice were given access to ethanol or water 4 h following removal from swim test cylinders. The non-stressed mice remained undisturbed in their home cage.

### Marble Burying Test

The marble burying test followed previously described procedures (38) and occurred when mice were 12–13 d withdrawn from CIE exposure and seven d withdrawn from FSS. Briefly, mice were placed in standard polycarbonate home cages with ~4 cm of fresh cob bedding for 2 h. They were returned to their home cage and 20 marbles were added to the test cage using a template. Mice were then placed in the test cage for 30 min and the number of buried marbles (>75%) was determined. For this experiment, mice were treated with CIE and FSS following standard methods but were not drinking ethanol in the home cage model.

### Novelty-Suppressed Feeding Test

Seven-eight days after the FSS (12-13 d withdrawn from CIE exposure), non-drinking mice were tested on the novelty-suppressed feeding test in a well-lit room, as described previously (7, 10). Briefly, mice were food deprived for 24 h and allowed to free feed for 2 h before a 22 h period of food deprivation.

Mice were allowed to habituate to the testing room for 1h before testing. At the start of NSFT, mice were then placed in the corner of an open field apparatus (40 x 40 x 40 cm) containing a single food pellet in the center and ~2 cm of fresh cob bedding on the floor. The latency to initiate food consumption was recorded and mice were immediately removed from the apparatus and placed in their home cages with a pre-weighed food pellet on the floor. Mice were allowed to consume the food pellet for 5 min. Vehicle (0.5% DMSO in 0.9% sterile saline; IP; 10 ml/kg dose volume), 1-EBIO (2.5–10 mg/kg; Tocris), or apamin (0.4 mg/kg; Sigma-Aldrich, St Louis, MO) was administered 30 min prior to behavioral testing.

### Bioinformatic analysis

Hypergeometric enrichment analysis was performed to determine if there was over-representation of significant correlations between the expression of each K^+^ channel gene in the BLA (represented by 91 probe sets targeting 88 unique K^+^ channel genes in the INIA Amygdala Cohort Affy MoGene 1.0 ST (Mar11) RMA Database) and 302 anxiety-like phenotypes from the BXD Published Phenotypes Database (7592 traits as of May, 2019) in male and female BXD RI strains of mice. All data sets used for this analysis are available in the GeneNetwork database (www.genenetwork.org; (39)). Spearman correlations between K^+^ channel expression and anxiety-like phenotypes that reached p < .01 were calculated for the analysis, and a Bonferroni correction was used to determine significance for the enrichment analysis. An eigentrait (the first principal component from principal components analysis) was computed in GeneNetwork from 40 behavioral traits. Only traits with sample size of 56 strains or greater were included. One strain (BXD11) that was an outlier was winsorized from 13.5 to 7.0. The *Kcnn3* eigentrait was mapped to regions of the genome containing gene variants that modulate trait expression using genome-wide efficient mixed model association (GEMMA; (40)) with the optional leave one chromosome out (LOCO) method in GeneNetwork2 (gn2.genenetwork.org; (41)). *Kcnn3* expression in the BLA (GeneNetwork Record ID 10493555) was correlated with phenotypes related to ethanol consumption.

### Long-access ethanol drinking

Before testing in the CIE-FSS drinking model, the ability of 1-ethyl-2-benzimidazolinone (1-EBIO) to reduce ethanol intake was established in a long-access 2-bottle choice paradigm. In this experiment, mice were given 22 h access to water or ethanol (15% v/v) until a stable baseline of consumption was established across five weeks of drinking. Vehicle or 1-EBIO (2.5, 5, and 10) was administered 30 minutes prior to ethanol availability, and intake levels were measured at 2, 4, and 22 h. Drug was administered once/week across three weeks in a counterbalanced within-subjects design. Non-cumulative drinking data at the different time points were used for analysis.

### Locomotor Testing

Locomotor activity was recorded in automated activity chambers (MED Associates, St. Albans, VT). C57BL/6J mice underwent a 10-min habituation session for three days followed by three daily 30-min test sessions. Thirty min prior to each habituation session, mice received an injection of vehicle. For test sessions, mice received either vehicle or 1-EBIO (2.5 or 10 mg/kg) in a counterbalanced within-subjects design.

### Statistical Analysis

A mixed data procedure (PROC MIXED) was used in the statistical software language SAS (SAS Institute Inc., Cary, NC) to analyze all drinking and behavioral data following our previous methods (37, 42). A within-subjects design was used when possible for these experiments to increase power and reduce variance associated with individual differences. In these repeated measures analyses, behavioral data were nested within mouse and further nested by time or session. This analytical approach was selected because of the capacity to handle unbalanced and complex repeated-measures data and the ability to model the variance and correlation structure of repeated-measures experimental designs (43). Tukey’s test was used for all post hoc analyses. All data are reported as mean ± SEM and statistical significance was established with p < 0.05.

## Results

### CIE and FSS effects on behavior

Consistent with previous studies (28–30), voluntary ethanol intake was increased in ethanol-dependent mice that were exposed to a 10 min FSS that occurred 4 h prior to the start of the drinking session (**Fig 1B**). Analysis revealed a significant interaction during the drinking session in test 2 (F(1,43) = 6.49, p = 0.0145; n = 11–12 mice/group), and post-hoc analysis showed that stressed, dependent mice drank more ethanol in 2 h than the mice in the other three treatment groups (p < 0.01).

During abstinence, mice were tested on two measures of anxiety-related behaviors. In the first cohort, mice were tested on the marble burying task. There was a significant interaction (F(1,39) = 5.91, p = 0.0197, n = 10–11 mice/group; **Fig 1C**) with post-hoc analysis revealing a difference between stressed mice in the air and CIE groups (p = 0.0031). A separate cohort of mice was tested on the NFST. There was a significant interaction (F(1,30) = 13.82, p = 0.0008; n = 7–10 mice/group; **Fig 1D**), and post-hoc tests revealed that stressed, ethanol-dependent mice had longer latencies to consume food in comparison with mice in non-stressed-air, stressed-air, and non-stressed-CIE groups (p < .01 vs all groups). Food consumption in the 5 min post-test was similar between all treatment groups (main effects: F(1,30) = 0.77; p = 0.387, and F(1,30) = 0.00, p ≥ 0.972; **Fig 1E**), suggesting that motivation to consume food in a familiar environment was not affected by CIE or FSS treatment, or their interaction.

### Genetic predictors of anxiety-related phenotypes

To identify K^+^ channel genes that are associated with anxiety-related phenotypes, we performed an unbiased targeted genetic screen using amygdala K^+^ channel transcript levels (91 probe sets) and phenotypic data (7500+) in BXD RI strains of mice. Of the published phenotypes, 301 were related to anxiety-like behaviors. Enrichment analysis identified 10 K^+^ channel genes that were over-represented with significant correlations with anxiety-like behaviors in BXD RI strains (**Fig 2A**). The top gene, *Kcnn3* (*Chr* 3), encodes a small-conductance, calcium-activated K^+^ (K_Ca_2.3 or SK3) channel that has been previously linked to alcohol-seeking behaviors (36). Further examination of *Kcnn3* expression in the amygdala revealed a strong eigentrait for anxiety-like and activity-based behaviors (Supplemental Table 1). Representative examples of correlations between BLA Kcnn3 expression and anxiety-like (i.e., avoidance) and activity in BXD RI strains are shown in **Fig 2B,C**. We next identified what loci control the variability in the Kcnn3-seeded network. Mapping analysis revealed that the eigentrait variance is polygenic, as there were multiple genomic regions on *Chr 1, 2*, and 4, but not on *Chr 3* near *Kcnn3* (at 89.52 Mb), containing gene variants significantly associated with eigentrait expression (**Fig 2D**). Interestingly, there were four SNPs in *Cacna1e* on *Chr 1* with significant association scores (**Fig 2D**). The R-type Ca^2+^ channel is encoded by *Cacna1e*, and R-type Ca^2+^ channels are functional coupled to K_Ca_2 channels (44, 45). These findings suggest that the *Kcnn3* ‘anxiety’ eigentrait is part of a complex phenotype involving genetic variation upstream (i.e., *Cacna1e*) of *Kcnn3*.

**Figure 2.**
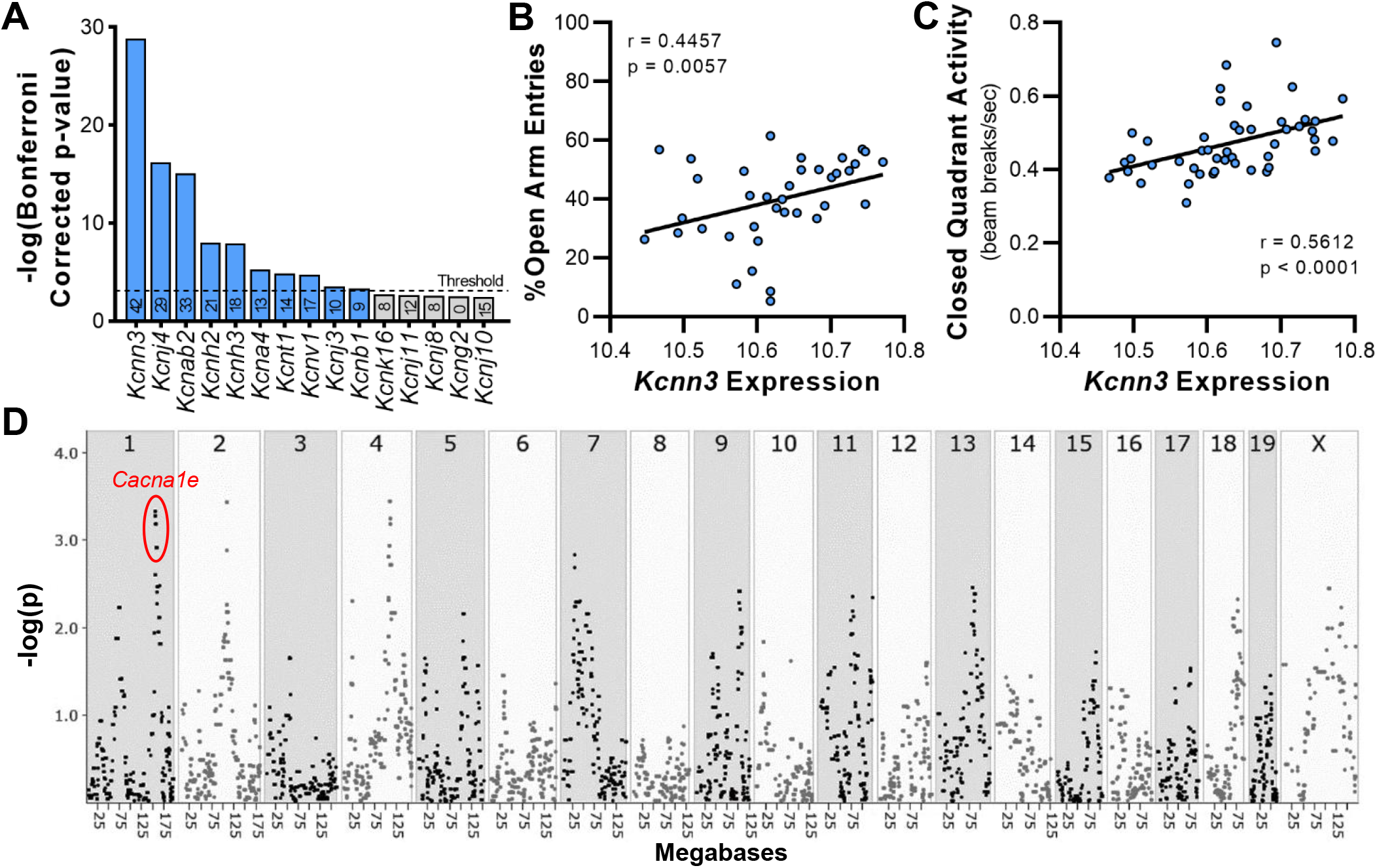
*Kcnn3* enrichment in anxiety-like and activity-based behaviors in BXD RI strains of mice. **(A)** Enrichment analysis of K^+^ channel genes in the BLA and anxiety-related phenotypes. **(B,C)** Correlations between BLA *Kcnn3* expression and open arm entries (record ID 11714) and closed quadrant activity (record ID 12354). **(D)** Mapping analysis of the Kcnn3-seeded eigentrait variance.

### BLA *Kcnn3* expression correlated with ethanol consumption

We next determined if *Kcnn3* expression in the BLA also correlated with voluntary drinking in BXD mice. As shown in **Table 1**, analysis revealed significant correlations between *Kcnn3* and ethanol consumption, the majority of which were negative (15 of 16 data sets). Significant correlations were observed in standard home cage drinking models, as well as the drinking in the dark model that produces binge-like consumption in C57BL/6J mice (46). Interestingly, seven of the 16 correlations identified between ethanol intake traits and *Kcnn3* expression are from mice that were exposed to a regiment of chronic mild stress. Representative correlations between BLA *Kcnn3* expression and ethanol drinking are shown in **Fig 3**. Given the strong relationship between amygdala *Kcnn3* and anxiety-like behaviors and voluntary drinking, a pharmacological approach was used to determine if increasing activity of K_Ca_2 channels with the positive modulator 1-EBIO would reduce anxiety-like behavior on the NSFT and excessive drinking in stressed, dependent mice.

**Figure 3.**
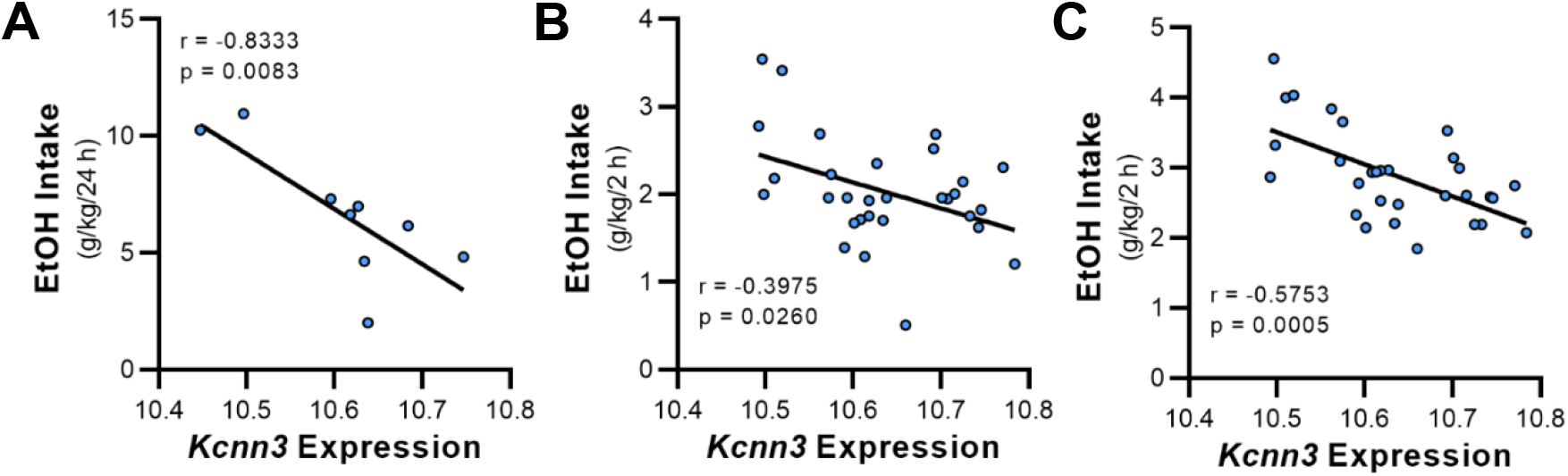
Negative correlations between BLA *Kcnn3* expression and home cage ethanol drinking in male and female BXD RI strains of mice. Significant correlations between BLA *Kcnn3* and consumption in **A)** a 24 h access model (3% v/v in 0.2% saccharin; record ID 10475); **B)** the baseline phase of the drinking in the dark model (20% v/v; record ID 20264), and **C)** DID drinking following chronic mild stress exposure (20% v/v; record ID 20288).

**Table 1.**
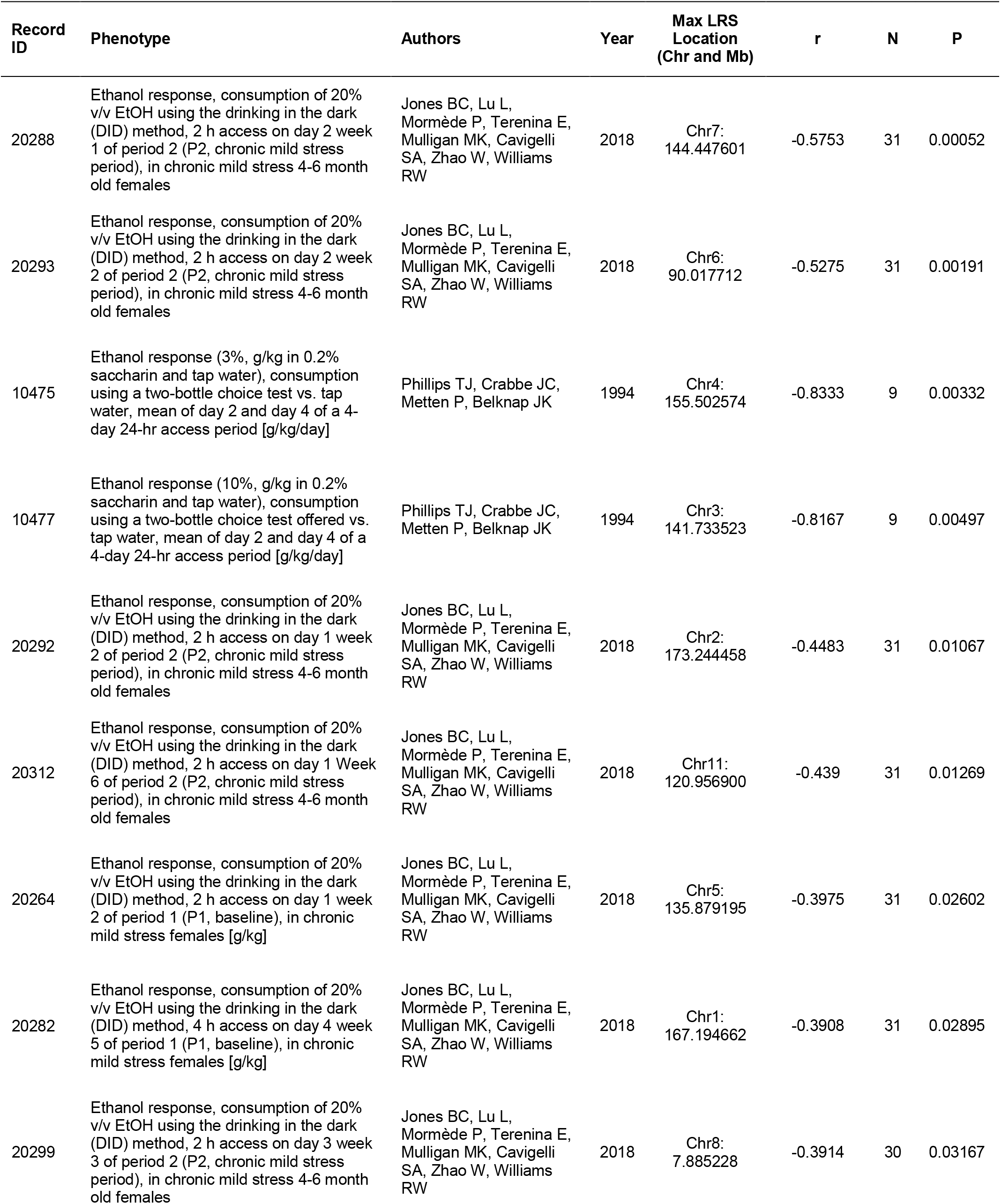
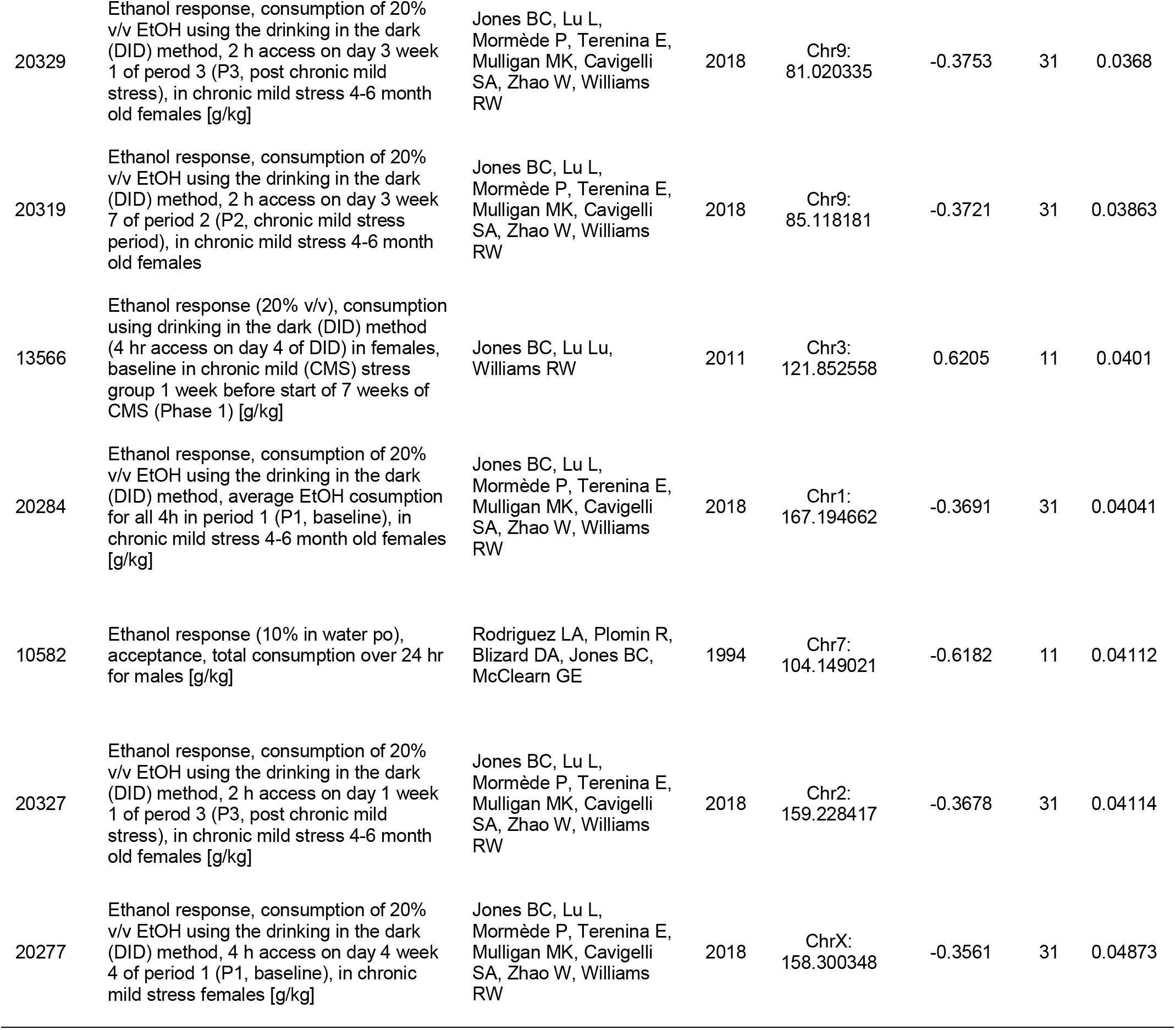
Correlations between *Kcnn3* expression in the BLA (INIA Amygdala Cohort Affy MoGene 1.0 ST (Mar11) RMA Database; Record ID 10493555) and voluntary ethanol drinking in male and female BXD RI strains of mice.

| Record ID | Phenotype | Authors | Year | Max LRS Location (Chr and Mb) | r | N | P |
| --- | --- | --- | --- | --- | --- | --- | --- |
| 20288 | Ethanol response, consumption of 20% v/v EtOH using the drinking in the dark (DID) method, 2 h access on day 2 week 1 of period 2 (P2, chronic mild stress period), in chronic mild stress 4-6 month old females | Jones BC, Lu L, Mormède P, Terenina E, Mulligan MK, Cavigelli SA, Zhao W, Williams RW | 2018 | Chr7: 144.447601 | -0.5753 | 31 | 0.00052 |
| 20293 | Ethanol response, consumption of 20% v/v EtOH using the drinking in the dark (DID) method, 2 h access on day 2 week 2 of period 2 (P2, chronic mild stress period), in chronic mild stress 4-6 month old females | Jones BC, Lu L, Mormède P, Terenina E, Mulligan MK, Cavigelli SA, Zhao W, Williams RW | 2018 | Chr6: 90.017712 | -0.5275 | 31 | 0.00191 |
| 10475 | Ethanol response (3%, g/kg in 0.2% saccharin and tap water), consumption using a two-bottle choice test vs. tap water, mean of day 2 and day 4 of a 4-day 24-hr access period [g/kg/day] | Phillips TJ, Crabbe JC, Metten P, Belknap JK | 1994 | Chr4: 155.502574 | -0.8333 | 9 | 0.00332 |
| 10477 | Ethanol response (10%, g/kg in 0.2% saccharin and tap water), consumption using a two-bottle choice test offered vs. tap water, mean of day 2 and day 4 of a 4-day 24-hr access period [g/kg/day] | Phillips TJ, Crabbe JC, Metten P, Belknap JK | 1994 | Chr3: 141.733523 | -0.8167 | 9 | 0.00497 |
| 20292 | Ethanol response, consumption of 20% v/v EtOH using the drinking in the dark (DID) method, 2 h access on day 1 week 2 of period 2 (P2, chronic mild stress period), in chronic mild stress 4-6 month old females | Jones BC, Lu L, Mormède P, Terenina E, Mulligan MK, Cavigelli SA, Zhao W, Williams RW | 2018 | Chr2: 173.244458 | -0.4483 | 31 | 0.01067 |
| 20312 | Ethanol response, consumption of 20% v/v EtOH using the drinking in the dark (DID) method, 2 h access on day 1 Week 6 of period 2 (P2, chronic mild stress period), in chronic mild stress 4-6 month old females | Jones BC, Lu L, Mormède P, Terenina E, Mulligan MK, Cavigelli SA, Zhao W, Williams RW | 2018 | Chr11: 120.956900 | -0.439 | 31 | 0.01269 |
| 20264 | Ethanol response, consumption of 20% v/v EtOH using the drinking in the dark (DID) method, 2 h access on day 1 week 2 of period 1 (P1, baseline), in chronic mild stress females [g/kg] | Jones BC, Lu L, Mormède P, Terenina E, Mulligan MK, Cavigelli SA, Zhao W, Williams RW | 2018 | Chr5: 135.879195 | -0.3975 | 31 | 0.02602 |
| 20282 | Ethanol response, consumption of 20% v/v EtOH using the drinking in the dark (DID) method, 4 h access on day 4 week 5 of period 1 (P1, baseline), in chronic mild stress females [g/kg] | Jones BC, Lu L, Mormède P, Terenina E, Mulligan MK, Cavigelli SA, Zhao W, Williams RW | 2018 | Chr1: 167.194662 | -0.3908 | 31 | 0.02895 |
| 20299 | Ethanol response, consumption of 20% v/v EtOH using the drinking in the dark (DID) method, 2 h access on day 3 week 3 of period 2 (P2, chronic mild stress period), in chronic mild stress 4-6 month old females | Jones BC, Lu L, Mormède P, Terenina E, Mulligan MK, Cavigelli SA, Zhao W, Williams RW | 2018 | Chr8: 7.885228 | -0.3914 | 30 | 0.03167 |
| 20329 | Ethanol response, consumption of 20% v/v EtOH using the drinking in the dark (DID) method, 2 h access on day 3 week 1 of period 3 (P3, post chronic mild stress), in chronic mild stress 4-6 month old females [g/kg] | Jones BC, Lu L, Mormède P, Terenina E, Mulligan MK, Cavigelli SA, Zhao W, Williams RW | 2018 | Chr9:<br>81.020335 | -0.3753 | 31 | 0.0368 |
| 20319 | Ethanol response, consumption of 20% v/v EtOH using the drinking in the dark (DID) method, 2 h access on day 3 week 7 of period 2 (P2, chronic mild stress period), in chronic mild stress 4-6 month old females | Jones BC, Lu L, Mormède P, Terenina E, Mulligan MK, Cavigelli SA, Zhao W, Williams RW | 2018 | Chr9:<br>85.118181 | -0.3721 | 31 | 0.03863 |
| 13566 | Ethanol response (20% v/v), consumption using drinking in the dark (DID) method (4 hr access on day 4 of DID) in females, baseline in chronic mild (CMS) stress group 1 week before start of 7 weeks of CMS (Phase 1) [g/kg] | Jones BC, Lu Lu, Williams RW | 2011 | Chr3:<br>121.852558 | 0.6205 | 11 | 0.0401 |
| 20284 | Ethanol response, consumption of 20% v/v EtOH using the drinking in the dark (DID) method, average EtOH consumption for all 4h in period 1 (P1, baseline), in chronic mild stress 4-6 month old females [g/kg] | Jones BC, Lu L, Mormède P, Terenina E, Mulligan MK, Cavigelli SA, Zhao W, Williams RW | 2018 | Chr1:<br>167.194662 | -0.3691 | 31 | 0.04041 |
| 10582 | Ethanol response (10% in water po), acceptance, total consumption over 24 hr for males [g/kg] | Rodriguez LA, Plomin R, Blizard DA, Jones BC, McClearn GE | 1994 | Chr7:<br>104.149021 | -0.6182 | 11 | 0.04112 |
| 20327 | Ethanol response, consumption of 20% v/v EtOH using the drinking in the dark (DID) method, 2 h access on day 1 week 1 of period 3 (P3, post chronic mild stress), in chronic mild stress 4-6 month old females [g/kg] | Jones BC, Lu L, Mormède P, Terenina E, Mulligan MK, Cavigelli SA, Zhao W, Williams RW | 2018 | Chr2:<br>159.228417 | -0.3678 | 31 | 0.04114 |
| 20277 | Ethanol response, consumption of 20% v/v EtOH using the drinking in the dark (DID) method, 4 h access on day 4 week 4 of period 1 (P1, baseline), in chronic mild stress females [g/kg] | Jones BC, Lu L, Mormède P, Terenina E, Mulligan MK, Cavigelli SA, Zhao W, Williams RW | 2018 | ChrX:<br>158.300348 | -0.3561 | 31 | 0.04873 |

### K_Ca_2 channels and aberrant behaviors in stressed, dependent mice

Non-ethanol drinking mice were treated with CIE and FSS as described previously and were then administered 1-EBIO (2.5 – 10 mg/kg) or vehicle 30 min prior to testing tested on the NSFT during withdrawal. A three-factor (air vs CIE, no FSS vs FSS, vehicle vs drug) analysis revealed a significant interaction (F(3, 123) = 4.25, p = 0.007, n = 7-11 mice/group; **Fig 4A**). There were also significant simple interactions for vehicle (F(1,123) = 13.35, p = 0.0004) and 2.5 mg/kg 1-EBIO (F(1,123) = 14.23, p = 0. 0003) but not 5 (F(1,123) = 0.25, p = 0.615) or 10 mg/kg (F(1,123) = 0.00, p = 0.946) doses of 1-EBIO. Post-hoc analyses revealed that stressed, dependent mice treated with vehicle or 2.5 mg/kg 1-EBIO had longer latencies to consume food during the NSFT compared with vehicle-treated mice in the three other groups, as well as compared with stressed, ethanol-dependent mice treated with 5 or 10 mg/kg 1-EBIO (all p values < 0.0001). Food consumption in the 5 min post-test was similar between all treatment groups (three factors: F(3,123) = 0.01; p = 0.391; **Fig 4B**). Simple interactions for treatment x group, treatment x dose, and group x dose were also not significant (P ≥ 0.612), suggesting that motivation to consume food in the home cage was not affected by 1-EBIO treatment in stressed, dependent mice.

**Figure 4.**
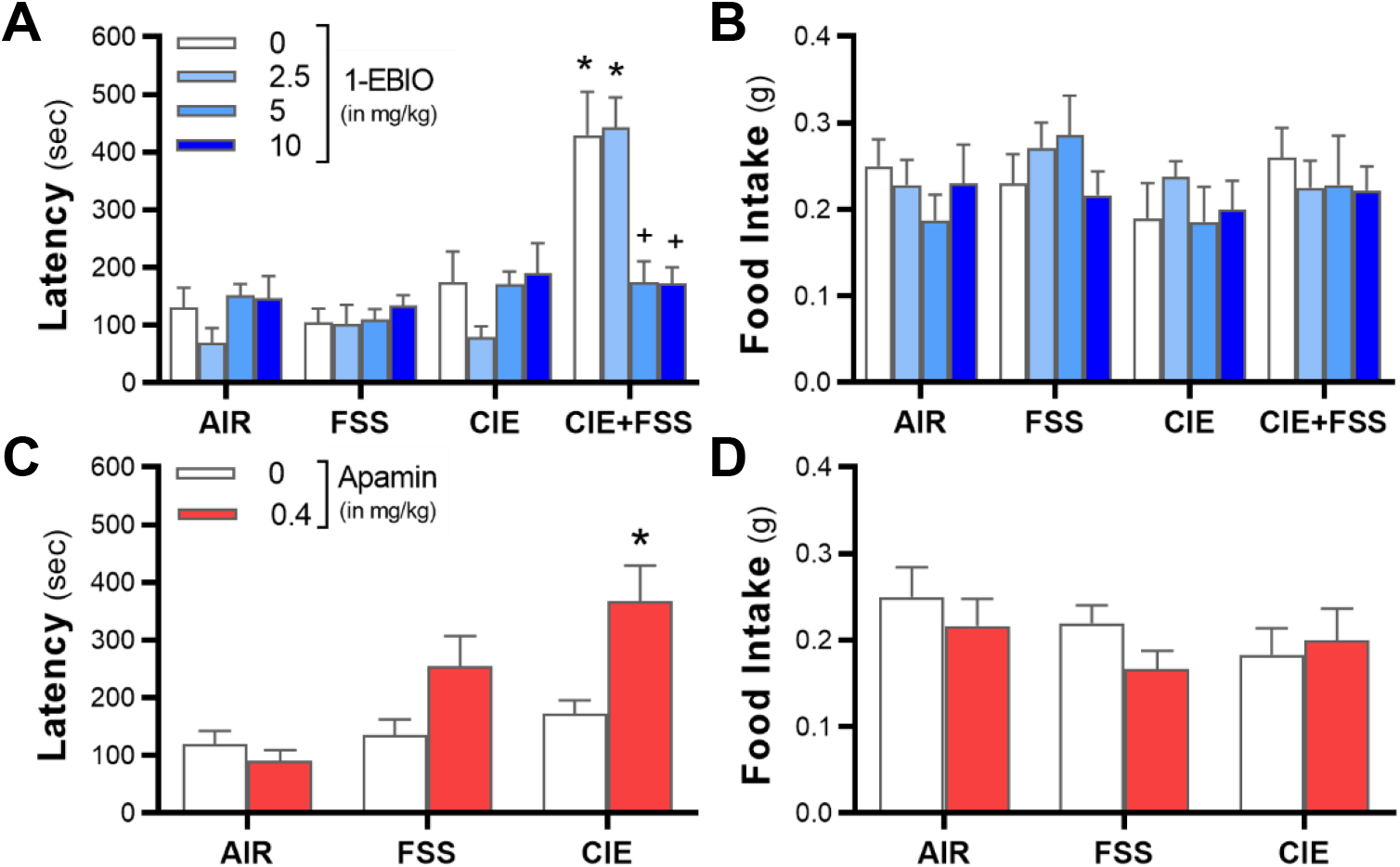
K_Ca_2 channels bidirectional regulate feeding behavior on the novelty-suppressed feeding test in stressed and ethanol dependent mice. **(A,B)** 1-EBIO significantly reduced latency to feed in stressed, dependent mice, without affecting post-test food consumption. **(C,D)** Apamin significantly increased latency to consume food in the NSFT in mice exposed to either FSS or CIE. Food intake in the home cage following the NSFT was not altered by apamin.

Next, we wanted to determine if K_Ca_2 channels have bidirectional control over feeding behavior on the NFST. We hypothesized that the allosteric inhibitor apamin would increase latency in mice with a history of FSS or CIE, similar to what was observed in stressed, dependent mice. There was a significant interaction with treatment and drug (F(2,28) = 4.03, p = 00291, n = 34 mice; **Fig 4C**). Post-hoc analysis showed that CIE exposed mice pretreated with 0.4 mg/kg apamin had longer latencies than vehicle-treated CIE exposed mice, as well as air controls treated with apamin or vehicle. Apamin administration nearly doubled the latency in FSS mice compared with air controls and FSS mice given vehicle, but this did not reach statistical significance. As before, food consumption in the 5 min post-test was similar between all treatment groups (F(2,28) = 0.73; p = 0.493; **Fig 4D**).

Because 1-EBIO reduced ethanol-seeking behaviors in rats (47, 48), we also wanted to test the ability of 1-EBIO to reduce excessive drinking in another species. C57BL/6J mice were given daily access to 22 h of ethanol (15%, v/v) or water. In this model, mice increased their drinking in weeks 3-5 in comparison with drinking amounts in the first week of access to ethanol (F(4,28) = 5.22, p = 0.003, n = 29 mice; posthoc, p < 0.05 vs week 1; **Fig 5A**). There was a significant interaction with time and drug dose (repeated measures, 2-factor design, F(6,80) = 3.66, p = 0.0029; **Fig 5B**). Post hoc analyses revealed that 10 mg/kg 1-EBIO significantly reduced drinking at the 2 h time point (p = 0.0151) in comparison with vehicle treatment.

**Figure 5.**
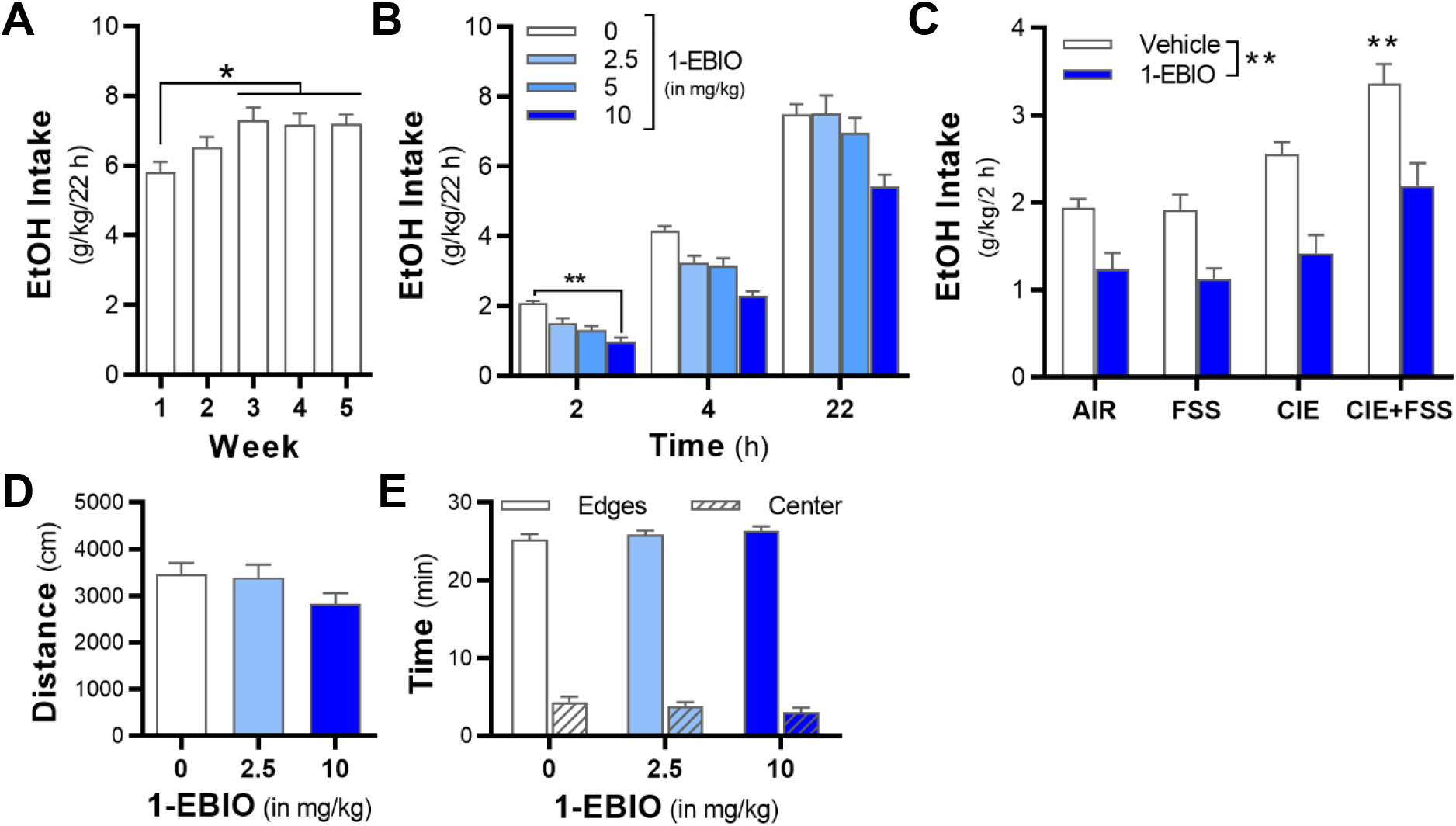
Positive modulation of K_Ca_2 channels reduced ethanol intake. **(A)** Mice escalate voluntary ethanol drinking when given 22 h of access to ethanol. 1-EBIO significantly reduced ethanol drinking in the **(B)** long- and **(C)** short-access home cage models regardless of treatment group. **(D,E)** 1-EBIO did not reduce overall distance traveled or time spent in the center or near the edges of an open field apparatus.

Mice exposed to the CIE and FSS paradigm were treated with vehicle and 10 mg/kg 1-EBIO in a within-subjects design across two drinking sessions during test 2. While the 3-way within-subjects interaction was not significant (F(1,42) = 0.02, p = 0.894; n = 11-12/group), there was a significant simple (stress by dependence) interaction in vehicle-treated mice (F(1,44) = 5.12, p = 0.029; **Fig 5C**). Consistent with results from the first cohort of mice, Bonferroni-corrected post-hoc analysis revealed that stressed, dependent mice consumed more alcohol than the other groups of mice (p < 0.003). All other simple interactions did not reach statistical significance (p values > 0.122). However, there were significant simple main effects of drug treatment when data were collapsed across the four treatment groups (F(1,42) ≥ 28.56, p < 0.0001), demonstrating that administration of 1-EBIO significantly reduced drinking in all groups of mice compared with drinking amounts after vehicle treatment. In an open field test, 1-EBIO did not affect overall locomotor behavior (repeated measures: F(2,28) = 2.38, p = 0.111, n = 18 mice; **Fig 5D**). While mice spent more time near the walls than the center of the open field apparatus (repeated measures: F(1,17) = 2158.17, p < 0.0001), there was no drug effect on time spent in center or edges (repeated measures: F(2,34) = 0.01, p = 0.986; **Fig 5E**).

## Discussion

Using a bioinformatics discovery tool, the *Kcnn3* gene was identified as the top K^+^ channel gene related to anxiety-like behaviors and was correlated with ethanol drinking in BXD RI strains. The expression of *Kcnn3* was also highly correlated with 40 anxiety- and activity-related behavioral traits, despite its moderate expression compared with the *Kcnn2* gene in the same K^+^ channel subfamily (49, 50). Genetic mapping of the eigentrait explaining the largest variance component of the correlated network of *Kcnn3* expression and behavioral traits revealed a complex and polygenic trait architecture with evidence of regulation by a locus on chromosome 1. This locus contained the gene encoding R-type Ca^2+^ channels that have known functional coupling with K_Ca_2 channels (44, 45). We then provided evidence for pharmacological validation of the *Kcnn3* gene product as a target to reduce stress-induced negative affective behaviors and stress-induced drinking in C57BL/6J mice. Specifically, we found that 1-EBIO reduced home cage ethanol drinking and reversed hypophagia druing the NSFT in stressed, dependent mice. We also found that blocking K_Ca_2 channels with systemic apamin administration in mice exposed to CIE exposure increased the latency on the NSFT, similar to what was observed in stressed, ethanol dependent mice. By demonstrating a role for *Kcnn3* in regulating excessive drinking and negative affective behaviors, these findings suggest that K_Ca_2 channels may be a key player in affective disorders and excessive drinking that are important contributors to the maintenance of AUD.

Previous evidence has revealed excessive ethanol intake and deficits in executive cognitive function in mice exposed to chronic intermittent ethanol and forced swim stress (28–31). Here, we found that this model of ethanol and stress co-exposure also produced anxiety-like behaviors in a novel environment without affecting food consumption in the home cage. This suggests that motivation to consume food was not different between treatment groups. Similar to the results from Rodberg and colleagues (2017), stressed, ethanol dependent mice did not bury more marbles than air controls, demonstrating that stress and ethanol co-exposure do not produce anxiety-like behaviors in all paradigms. In contrast to existing reports (38, 51, 52), ethanol exposure in the absence of repeated stress did not increase the number of marbles buried or the latency on the NSFT. The reason for the discrepancy is unknown but may relate to the relatively short duration of ethanol exposure or the time of testing relative to the last ethanol exposure. In our study, mice were exposed to two cycles of CIE exposure with a week in between and were tested on the behavioral assays two weeks after removal from the ethanol vapor chambers. Together, these findings suggest that the development of anxiety-like behaviors and cognitive deficits in stressed and ethanol dependent mice is complex and may depend on the model and time since last exposure.

Our pharmacological studies using a K_Ca_2 channel positive modulator demonstrated that we could prevent the hypophagia behavior observed on the NFST in the stressed, dependent mice without affecting the latency in the other treatment groups. Similarly, 1-EBIO reduced the latency to feed on the NFST in mice exposed to acute foot shock stress (53). Complementary to these findings, we found that blocking K_Ca_2 channel activity increased latency on the NFST in ethanol dependent mice, suggesting a potential interaction between stress and ethanol on K_Ca_2 channel function. Previous evidence has demonstrated that adolescent ethanol exposure increased anxiety levels and reduced K_Ca_2.3 channel expression in the nucleus accumbens shell (54). K_Ca_2 channels are known regulators of the medium afterhyperpolarization (mAHP) (55), and the amplitude of the mAHP was reduced in the adolescent exposed rodents, which is consistent with a loss of K_Ca_2 channel currents (54). In these animals, 1-EBIO microinfusion into the accumbens shell attenuated anxiety-like behavior in adolescent exposed rats. In addition, chronic ethanol exposure in rodents reduced K_Ca_2 channel function and/or expression in the accumbens core, orbitofrontal cortex, medial prefrontal cortex, ventral hippocampus, and ventral tegmental area (36, 47, 56–58), and *KCNN2* expression was reduced in postmortem BLA and frontal cortex of individuals with AUD (59). In a recent study, heavy ethanol drinking and ethanol dependence hypermethylated exon 1A of *KCNN3* and increased expression of apamin-insensitive and dominant-negative isoforms of K_Ca_2.3 channels in the nucleus accumbens of mice and monkeys (60). Using the same bioinformatic analysis as was used for the BLA, accumbens expression of *Kcnn2* or *Kcnn3* was not enriched with correlations for anxiety-related phenotypes in BXD RI strains (data not show).

In a model for the study of anxiety-like behavior, K_Ca_2.3 channel expression was upregulated in the dorsal raphe nucleus of socially isolated adolescent mice, and blocking K_Ca_2 channels partially reversed the increased latency reported on the NFST and normalized depressive-like behaviors on the tail suspension and sucrose preference tests in single-housed mice (61). In contrast, socially-isolated adolescent rats showed reduced BLA expression of K_Ca_2.2 and K_Ca_2.3 channels, and 1-EBIO reduced anxiety-like behaviors when microinfused into the BLA and bath application normalized hyperexcitability in these BLA pyramidal neurons (62). In adult mice, chronic restraint stress increased anxiety-like behaviors and reduced K_Ca_2.2 channel currents in BLA projection neurons, and K_Ca_2.2 channel overexpression in ventral hippocampal-projecting BLA neurons prevented anxiety-like behaviors in stressed mice (23). Although the relationship between ethanol, stress, and K_Ca_2 channels is complex and depends on brain region and age of stress or ethanol exposure, our findings add to a growing literature that K_Ca_2 channels are a crucial ion channel family that may be a possible pharmacogenetic target for treating comorbid mood and alcohol use disorders.

In addition to reducing hypophagia in stressed, dependent mice, we also showed that 1-EBIO attenuated ethanol drinking in long- and short-access models. In the 22 h drinking model, the reduction in drinking by 1-EBIO was only observed at 2 h time point, although intake remained below values in vehicle treated mice at the later time points. In the stress-ethanol co-exposure model, 1-EBIO decreased drinking in all groups, including the control, non-stressed mice. While the half-life of 1-EBIO is not reported in the literature, the relatively short-lived reduction in ethanol intake suggests that 1-EBIO may be rapidly metabolized in adult mice. These findings with 1-EBIO are consistent with previous evidence for reductions in drinking using K_Ca_2 channel positive modulators across multiple models of ethanol voluntary intake and seeking behaviors in mice and rats (47, 48, 63). In addition, we previously reported that *Kcnn* gene expression in the nucleus accumbens of BXD RI strains negatively correlated with multiple models of voluntary drinking, including binge-like consumption and CIE-induced escalation of drinking (35, 36). This suggests that K_Ca_2 channels are an effective target to reduce moderate and excessive amounts of ethanol, especially since effective doses of 1-EBIO do not appear to have adverse effects on locomotor activity.

In summary, the bioinformatics and pharmacological results demonstrate a role for K_Ca_2 channels in mediating anxiety-like behaviors and excessive ethanol drinking in mice. Recently, a small-scale clinical trial in social and moderate drinkers (ClinicalTrials.gov record number: NCT01342341) was completed with chlorzoxazone, an FDA-approved K_Ca_2 channel positive modulator (64) that is prescribed as a skeletal muscle relaxant (65). Chlorzoxazone was well tolerated, but it did not reduce the number of drinks across the two weeks of treatment. The reasons for the discrepancy between the rodent studies and clinical trial are unclear but may relate to chlorzoxazone’s short half-life, low EC_50_ for K_Ca_2 channels, off-target actions, conservative dosing approach, or population of drinkers enrolled in the trial. Despite the failed trial, K_Ca_2 channels remain a viable target to treat individuals with AUD and comorbid mood disorders.

## Acknowledgements

These studies were supported by NIH grants AA020930 (PJM), AA023288 (PJM), AA025110 (JAR), AA014095 (HCB), AA020929 (MFL), AA013499 (RWW), AA016662 (RWW), and AA017590 (RWW).

---

**Supplemental Table 1.**
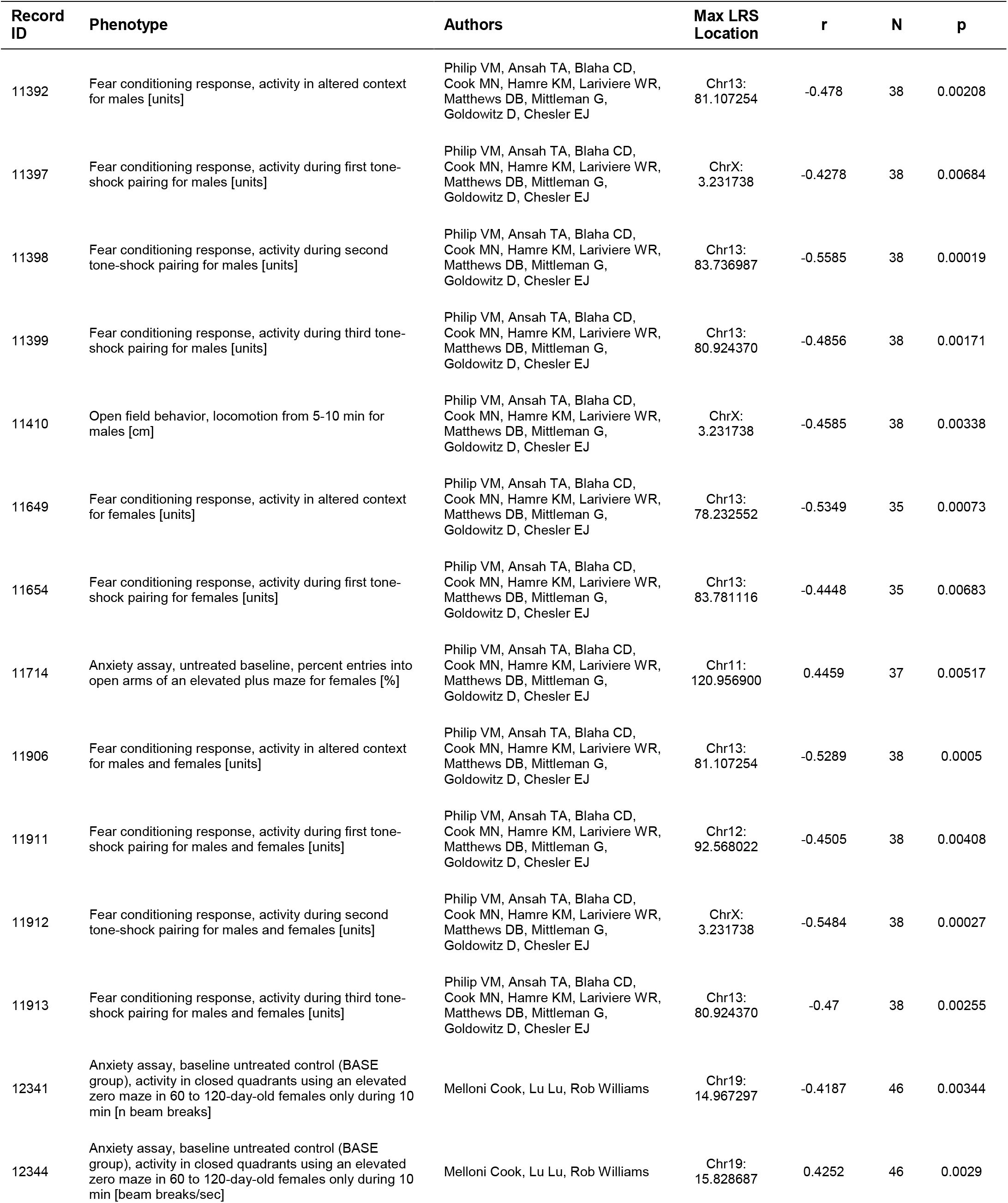
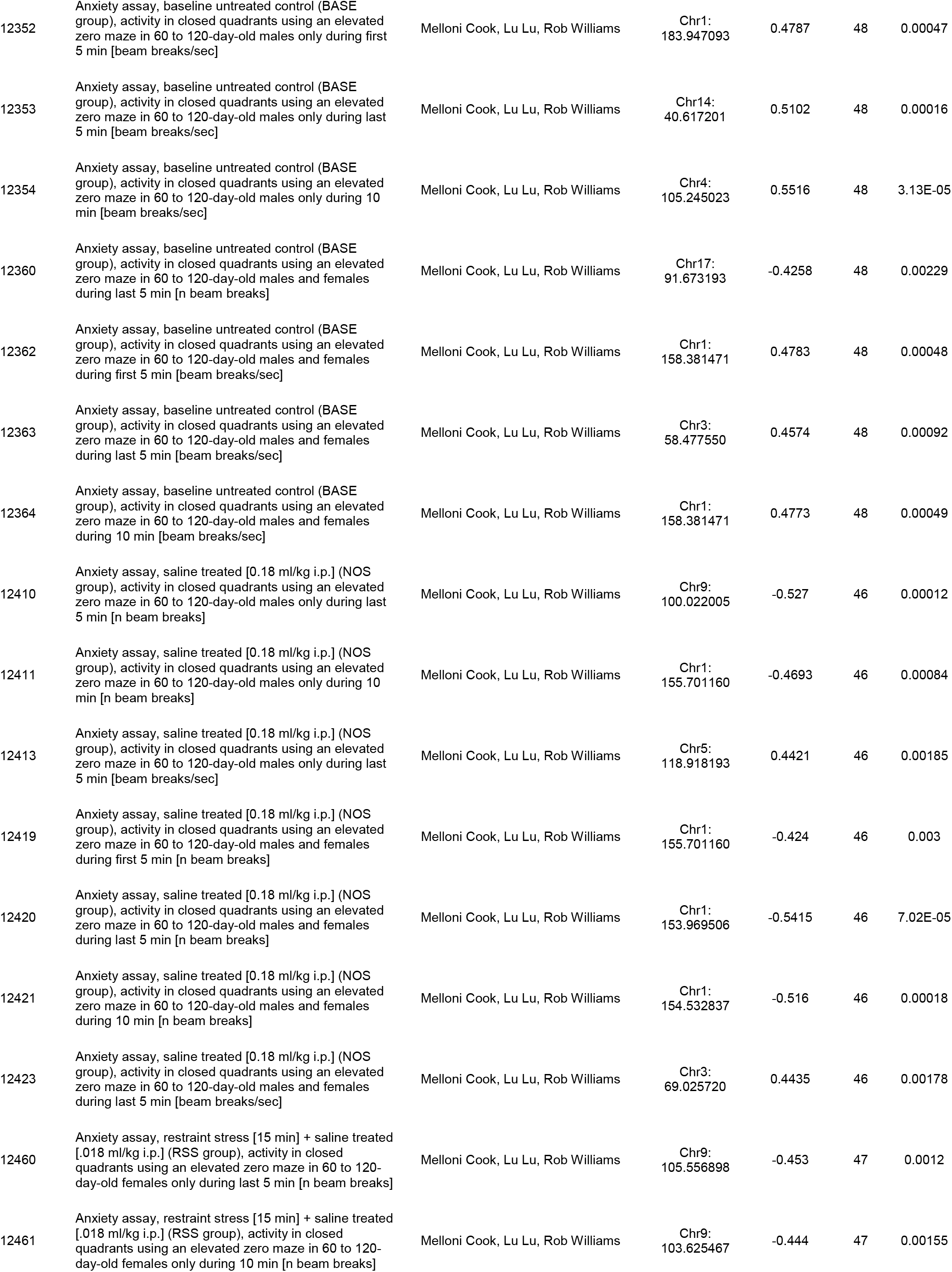
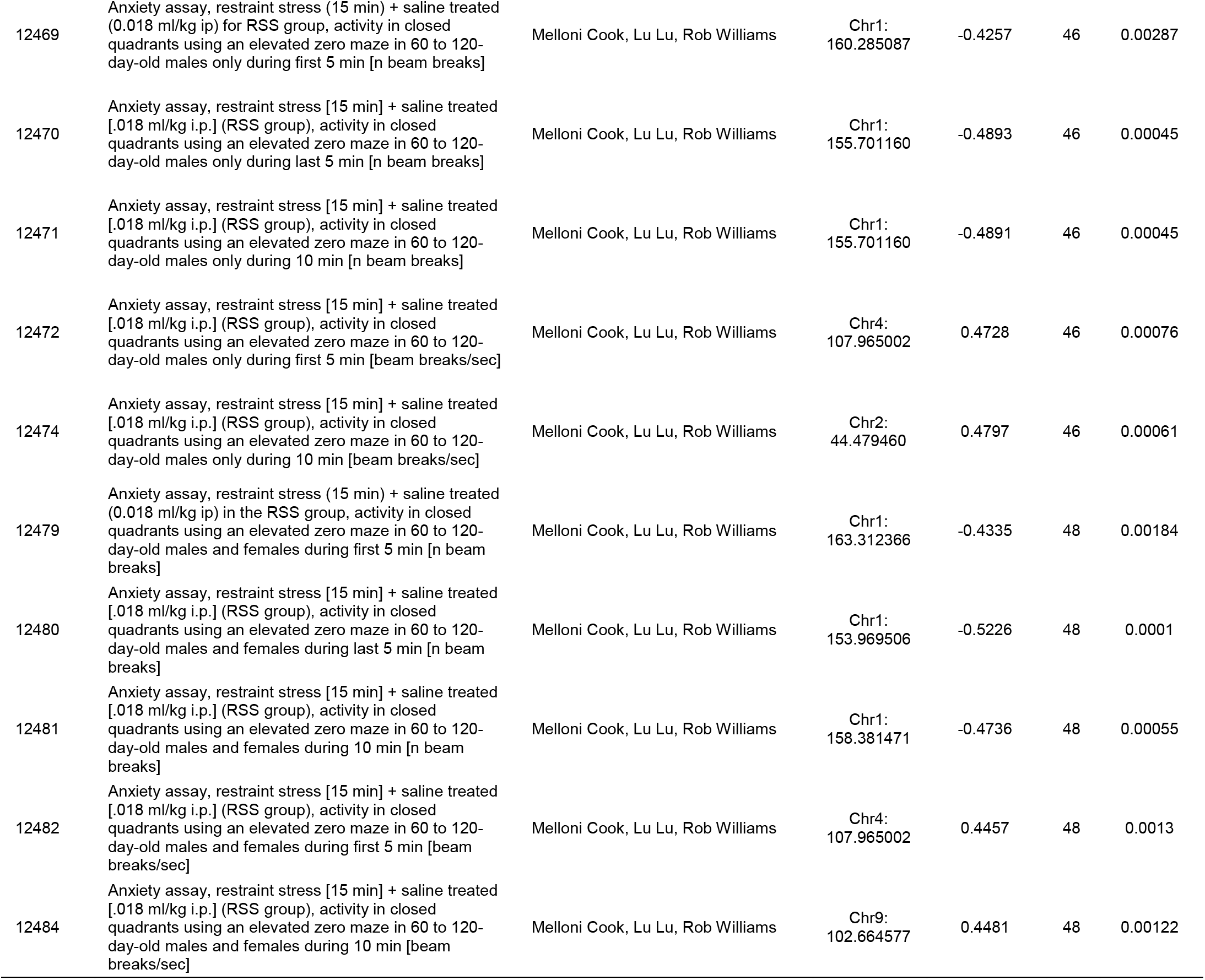
Anxiety-like behavior and activity eigentrait computed from 40 behavioral traits with sample size of 56 strains or greater (GeneNetwork Record ID 18794). Note that BXD11 was an outlier and data were winsorised from 13.459 to 7.0. All data are publically available on genenetwork.org.

